# Multi-reactive hydrogel nanovials for temporal control of secretion capture from antibody-secreting cells

**DOI:** 10.1101/2024.12.10.627574

**Authors:** Michael Mellody, Yuta Nakagawa, Richard James, Dino Di Carlo

**Affiliations:** Department of Bioengineering, University of California, Los Angeles; Center for Immunity and Immunotherapies, Seattle Children’s Research Institute; Department of Pediatrics and Pharmacology, University of Washington, Seattle; California Nanosystems Institute, University of California, Los Angeles; Jonsson Comprehensive Cancer Center, University of California, Los Angeles

## Abstract

Antibody discovery can benefit from techniques to screen antibody-secreting cells (ASCs) at scale for the binding and functionality of a diverse set of secreted antibodies. Previously, we demonstrated the use of cavity-containing hydrogel microparticles (nanovials) coated with a single affinity agent, biotin, to capture and identify ASCs secreting antibodies against a recombinant antigen bound to the nanovial through biotin-streptavidin linkages. However, rapidly secreted antibodies from unbound cells or cells in adjacent nanovials can cause crosstalk leading to background signal. Earlier efforts address this by localizing capture sites to the nanovial cavity, emulsifying nanovials, or short secretion times to limit secreted antibodies from binding to neighboring nanovials. Here, we demonstrate a method to functionalize nanovials with moieties that impart orthogonal reactivity, enabling conjugation of cell capture antibodies and antigens at different times. We show that by using a strained alkyne moiety to attach cell-capture antibodies via click chemistry to nanovials, we can capture cells and subsequently quantify secretions via biotin-streptavidin linkages. By delaying the loading of antigens onto the nanovials until after cell capture, we were able to ensure high purity (>95%) isolation of hybridoma secreting an antigen-specific antibody in a background of other hybridoma. This approach allows tight temporal control of the secretion measurement, which is independent of the cell loading time, and requires less convective transfer steps. Click chemistry-based coupling further improved cell loading into nanovials by 58% compared to biotin-streptavidin-biotin coupling and caused no reduction in cell viability. We demonstrate an implementation of this system to improve antigen-specific hybridoma screening, yielding an 8-fold improvement in hybridoma enrichment while maintaining a similar workflow complexity. Hybridomas on nanovials sorted into well plates regrew into colonies following sorting using standard fluorescence-activated cell sorting and maintained secretion of antigen-specific antibodies with high purity (∼90%), as validated via standard enzyme-linked immunosorbent assays. This lab-on-a-particle approach can be applied more generally to decouple cell loading, treatment, or activation, from secretion measurements for single-cell functional assays.

## Introduction

Monoclonal antibodies (mAbs) are vital to modern medicine and biomedical research due to their ability to specifically target and bind unique antigen epitopes.^1^ This specific binding enables more effective treatments with fewer side effects for a wide range of conditions, including several types of cancer, autoimmune disease, viral infection, and others.^2^ mAbs are also used in diagnostic applications to detect analytes present in body fluids or other samples and in biological research to measure levels of analytes in bulk samples (e.g. western blots) or at the single-cell level (e.g. flow cytometry), among other applications.^3^

Antibody discovery is based on a system of generating a diverse pool of mAb clones that target a specific antigen. These pools can be generated from immunization campaigns which are performed using antibody-secreting hybridomas, memory B cells, or plasma cells. The first to be developed, hybridoma technology, was invented in 1975 and is well-documented.^4,5^ Briefly, a mouse (or another suitable host) is immunized with an antigen the desired antibody is to be raised against. B cells are harvested from the spleen of the animal and fused to myeloma cells, creating a large population of long-lived, proliferating, and secreting hybridomas.^6^ Memory cells are long-lived B cells that have undergone affinity maturation and are specific to a previously encountered antigen. These have been used to develop therapies for infectious diseases such as HIV and SARS-CoV-2.^7,8^ Innovation in primary B cell culture has enabled direct sequencing of plasma cell antibodies without hybridoma generation.^9–11^

All of these campaigns can generate heterogeneous populations of hundreds of thousands to millions of unique antibody-secreting cells (ASCs).^12^ These ASCs are then screened based on the binding or other properties of the antibodies they secrete, usually using enzyme-linked immunosorbent assays (ELISAs) or other measurements of cell binding. However, screening this full diversity is not feasible with traditional well-plate-based screening systems due to their limited throughput. Growing clonal populations of hybridomas in multi-well plates also requires weeks to accumulate enough secreted antibodies for characterization. Pooling many clones in shared wells can help improve throughput, but can lead to enrichment of fast-growing, low affinity clones. Consequently, only a fraction of the potential antibody sequence space is usually surveyed, resulting in missed antibody clones with potentially more advantageous properties. To screen a higher depth of the antibody repertoire, there is a need for highly scalable single-cell tools to isolate and enrich individual ASCs based on the properties of secreted antibodies, without the time or material cost for regrowth.

Numerous single-cell screening approaches have been explored to isolate and enrich ASCs, including microwell arrays, optofluidic systems, and microfluidic droplets. Microwell arrays are used to isolate individual ASCs in separate wells, where fluorescence or enzymatic detection of secreted antibodies can be performed.^13^ Optofluidic instruments like the Beacon^TM^ (Berkeley Lights, now Bruker Cellular Analysis) are automated systems that similarly discretize cells into individual compartments and enable continuous monitoring of cell behavior via imaging and functional assays.^14,15^ However, drawbacks in both platforms include their limited throughput of ∼10^3^ cells per chip and high cost to operate. As an alternative, microfluidic droplet systems can be used to create millions of individual compartments for single-cell antibody secretion assays. Secreted antibodies can be rapidly accumulated within droplets and either fluorescence resonance transfer assays or beads coated with antigen are often used for localization and detection.^16^ Despite this, microfluidic droplets are still inaccessible to most researchers due to the complexity needed to generate, analyze, and sort droplets in a robust manner.^17,18^ Further, reactions in droplets are difficult to wash, leading to background signal that accompanies unbound detection reagents in the homogeneous droplet assay format. The ideal tool would compartmentalize hundreds of thousands to millions of ASCs and be compatible with standard fluorescence activated cell sorting (FACS) systems.

We have previously described a cavity-containing hydrogel microparticle system (nanovials) to load single ASCs and capture antigen-specific antibodies on the particle surface.^19,20^ Nanovials have also been applied to discover new antibodies from plasma cells extracted from immunized mice, by capturing mAbs that bind antigens conjugated to nanovials or bind to antigens expressed on the surface of target cells co-loaded onto nanovials.^21^ Importantly, nanovials are compatible with standard laboratory techniques and do not require extraneous microfluidic equipment after they are fabricated. The latest implementation of nanovials is fabricated using droplets comprising of an aqueous two-phase system that phase separates into a polyethylene glycol (PEG)-rich outer region and gelatin-rich inner region to form crosslinked PEG-based hydrogels with a thin layer of gelatin in the cavity. The open-faced nanovial cavity holds picoliter-scale volumes and can be functionalized with biotin, antibodies, or antigens via N-hydroxysuccinimide (NHS)-ester chemistries. We leverage local functionalization within the cavity to capture cells and specific secretions from those cells. After labeling the nanovial cavities with antibodies to capture cells and antigens to capture secreted antibodies, ASCs are loaded onto nanovials. Following incubation, unbound ASCs are removed and the remaining cell-loaded nanovials are incubated for secretion accumulation. These are stained with a fluorescent detection antibody and sorted via FACS, enabling screening of millions of clones in a matter of hours.

However, during cell loading and mixing steps, antibodies secreted from captured and unbound ASCs can lead to background signal.^19,20^ Empty nanovials and nanovials containing poorly-secreting or low-affinity ASCs may capture some of these secretions from unbound cells, leading to reduced signal to noise and reduced purity of the final enriched pool. This is driven by the fact that nanovials are already decorated with biotinylated secretion-capture proteins during the initial ASC loading step. During this step both biotinylated secretion-capture and cell-capture proteins are added simultaneously to control the relative amounts of both proteins so that the secretion-capture proteins are not out-competed for available streptavidin sites. This results in little temporal control of when secretions can be captured from cells and higher rates of crosstalk from cells secreting during cell loading steps. Using gelatin extracellular matrix on the nanovials to load cells prior to adding secretion capture moieties, localizing capture sites on the gelatin, reducing secretion times, emulsifying nanovials, and adding blocking antibodies can localize secretions or act as a sink for unbound antibodies but these approaches have tradeoffs with more complex workflows, limitations to adherent cells, and/or reduced signal.^19,20,22,23^ New approaches for temporally separating secretion capture from cell capture on nanovials would be advantageous to further improve assay flexibility, purity of isolated clones, and to develop new assay formats that provide temporal information about secretions.

Here, we modify nanovials to introduce orthogonal chemistries that allow us to decouple cell loading and antigen-loading onto nanovials in time, which we hypothesized could reduce crosstalk and ultimately improve hybridoma selection and mAb discovery (Fig 1). Following microfluidic fabrication of nanovials (Fig 1A) upstream of the ASC assay, nanovials are decorated with NHS-dibenzocyclooctyne (DiBO) groups, and cell-capture antibodies are functionalized with azide groups (Fig 1B). The azido-modified antibody reacts with DiBO through a strain-promoted azide-alkyne cycloaddition to covalently conjugate the antibody to the nanovial cavity. We demonstrate that by using click chemistry moieties in parallel to biotin-streptavidin, it is possible to temporally separate cell loading from secretion accumulation on nanovials. This technique increases specific signal from ASCs to >40-fold the background signal on empty nanovials and results in enriched cell purity via FACS sorting on commercial instruments for scenarios where crosstalk can dominate with workflows applied to uniformly functionalized nanovials (Fig 1C). Importantly, this approach requires only small adjustments to the original nanovial workflow, making it well-adapted for other cell and assay types (Fig 1D). This innovation further strengthens the nanovial platform for antibody development and removes limitations in studying temporally-dependent secretion patterns for numerous other applications.

**Figure 1.**
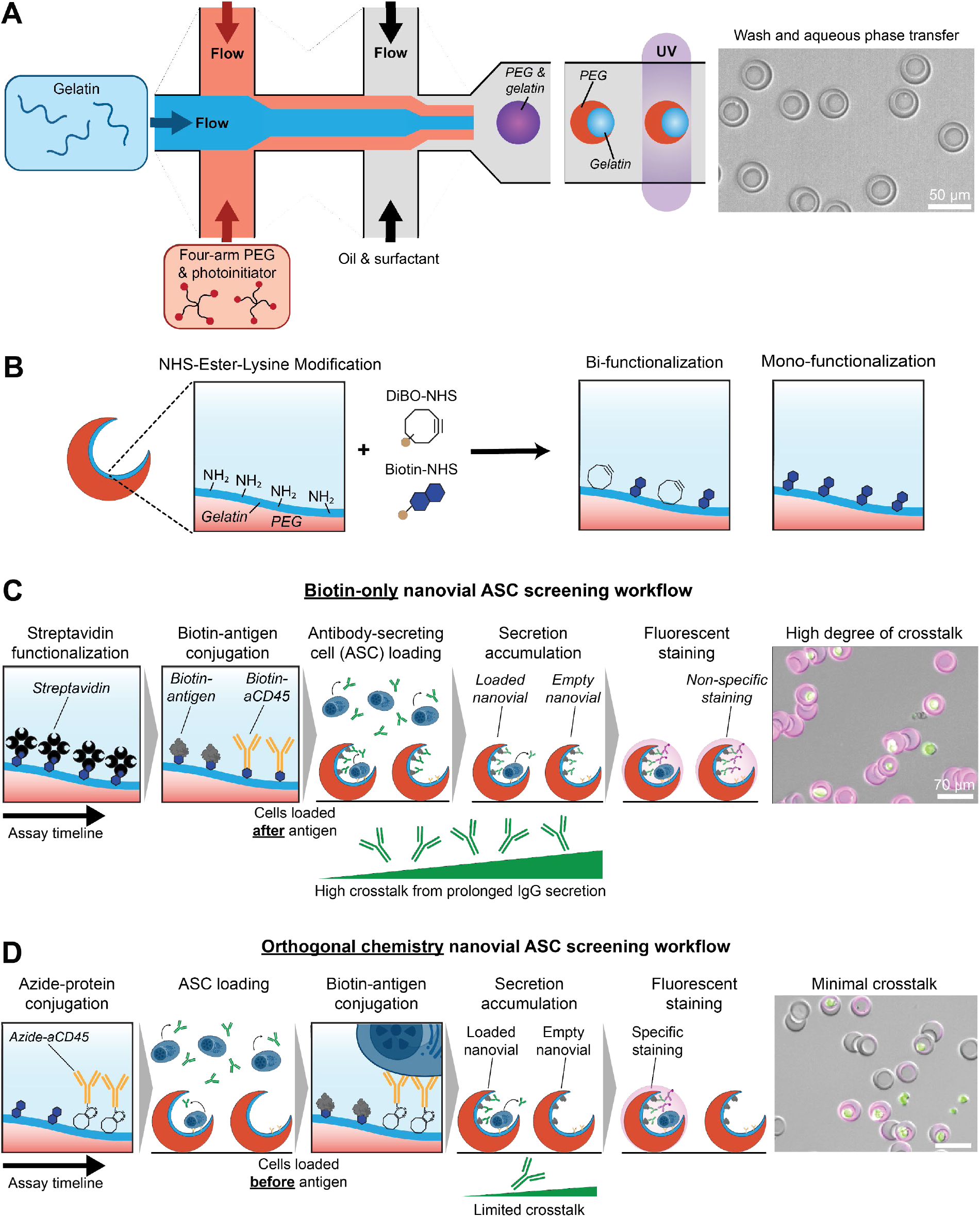
Multi-reactive nanovial workflow for enrichment of antigen-specific antibody secretion capture. A) Nanovials are fabricated using an aqueous two-phase system, resulting in PEG-based particles containing a thin layer of gelatin embedded within the inner cavity. B) NHS-ester chemical moieties are attached to the lysine groups on the gelatin layer via amide bonds. Biotin and dibenzocyclooctyne (DiBO) groups can be conjugated in parallel or individually. C) In the biotin-only workflow, cell capture and secretion capture proteins are conjugated upstream of cell loading, which results in increased non-specific secretion capture and crosstalk. D) By using DiBO-azide (click chemistry) as an orthogonal chemistry, cell capture antibodies can be conjugated prior to secretion capture proteins, which minimizes non-specific binding.

## Results

### Generating nanovials with orthogonally-reactive surface moieties

Nanovials are fabricated using microfluidic droplet generation of water-in-oil droplets comprising a PEG-gelatin aqueous two-phase system which rapidly produces millions of PEG-based particles containing a lysine-rich inner layer of gelatin (Fig 1A). After washing away excess oil and surfactant, nanovials are de-emulsified and in aqueous phase for every procedure beyond this point. Nanovials can then be chemically modified for various cellular assays. We have leveraged NHS ester-amine conjugation chemistries to attach biotin molecules to lysine groups in the gelatin layer (Fig 1B). The development of a multi-reactive nanovial surface is contingent on two or more chemical moieties binding to the nanovial cavity and being non-reactive to each other. We conjugated nanovials with different ratios of NHS-DiBO and NHS-biotin, then attached Alexa

Fluor 647 streptavidin and FITC azide to assess surface modification and orthogonal reactivity (Fig 2A). Fluorescence microscopy revealed a predictable trend where increased NHS-DiBO concentrations corresponded to higher FITC signals and higher biotin concentrations yielded increased Alexa Fluor 647 fluorescence. We incubated nanovials with a range of NHS-DiBO concentrations both with and without 10 mM NHS-biotin (Fig 2B & 2C). Flow cytometry analysis revealed competitive binding of DiBO and biotin onto the gelatin cavity. In the absence of biotin, DiBO levels were elevated over 6-fold compared to co-conjugation with 10 mM biotin. As expected, the presence of DiBO (1 mM) reduced the relative abundance of biotin by almost 4-fold. We also found that DiBO-NHS with a PEG spacer led to improved surface decoration compared to a shorter sulfone spacer (SI Fig 1A). All further multi-reactive nanovials were conjugated with 1 mM DiBO-PEG-NHS and 10 mM biotin-NHS. We further showed that nanovials functionalized with DiBO and biotin and stored at 4°C remained stably conjugated for at least two weeks post-functionalization (SI Fig 1B). The long-term stability of orthogonal chemistry groups on nanovials demonstrates the potential for this system in multi-day cellular assays where a consistent nanovial surface profile is essential. We next validated that azido-modified antibodies could be conjugated to the multi-reactive nanovial surface. We found that after functionalization, anti-mouse CD45 antibodies could be conjugated to the nanovial surface in a dose dependent fashion for the same incubation period as its biotinylated version (Fig 2D). In addition, we confirmed that azido-modified antibodies were restricted to the nanovial cavity. Conjugation was detected using an Alexa Fluor 647 anti-IgG antibody and image analysis was performed using ImageJ.

**Figure 2.**
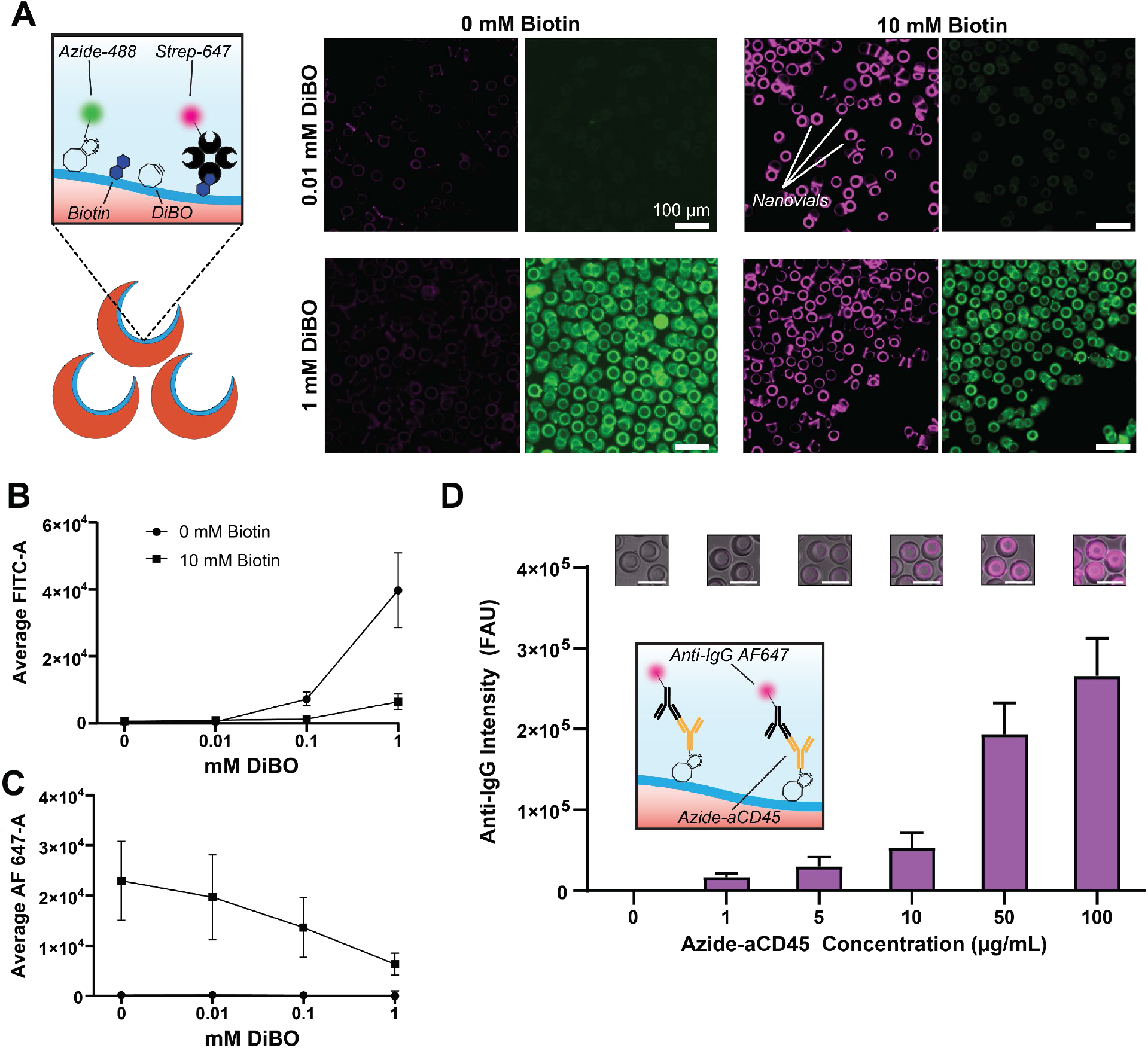
Characterization of multi-reactive nanovials. A) Schematic of experiment and fluorescence microscopy images of nanovials coated with biotin and/or DiBO and reacted with azide-FITC (green) and streptavidin-Alexa Fluor 647 (magenta). Scale bars are 100 microns. Left images are in Cy5 emission channel and right images are in FITC emission channel. B) Flow cytometry measurements of FITC and Alexa Fluor 647 signal area of nanovials coated with biotin and/or DiBO. C) Fluorescence microscopy and image analysis of DiBO-modified nanovials conjugated with azido-modified IgG antibody (aCD45) and detected with Alexa Fluor 647 anti-IgG. Scale bars are 50 microns.

### Loading cells onto nanovials using antibodies modified to link to either surface chemistry

We then sought to assess the functionality of azido-modified antibodies for cell loading into nanovials. To do this, we compared the loading efficiency of HyHEL5 hybridoma cells using azido-modified anti-CD45 antibodies linked to the nanovials through DiBO versus biotinylated anti-CD45 antibodies bound to streptavidin coated nanovials (Fig 3A). HyHEL5 hybridoma cells have high levels of CD45 on their membranes and can be stably loaded into nanovials through antibody binding. Loading efficiency was defined as the fraction of nanovials containing at least one cell when mixed with cells at a ratio of 180,000 nanovials to 300,000 cells. Staining cells with calcein AM enabled high-throughput screening of tens of thousands of nanovials by flow cytometry in a matter of minutes to obtain robust loading statistics (Fig 3B). Nanovials functionalized with azide-conjugated antibodies demonstrated superior binding across every tested concentration, with a maximum efficiency of 37.9% at 150 μg/mL. Nanovials functionalized with biotinylated anti-CD45 antibodies reached loading saturation at 50 μg/mL with a loading efficiency of 21.0% (Fig 3C). The improved loading of azide-conjugated nanovials may be due to the covalent nature of the click chemistry linkage and reduction of other steric interference from large proteins such as may be present with biotin-streptavidin linkages. These results highlight the potential of click chemistry-based nanovials to improve cell loading rates and pan large pools of cells for rare clones with less cell loss.

**Figure 3.**
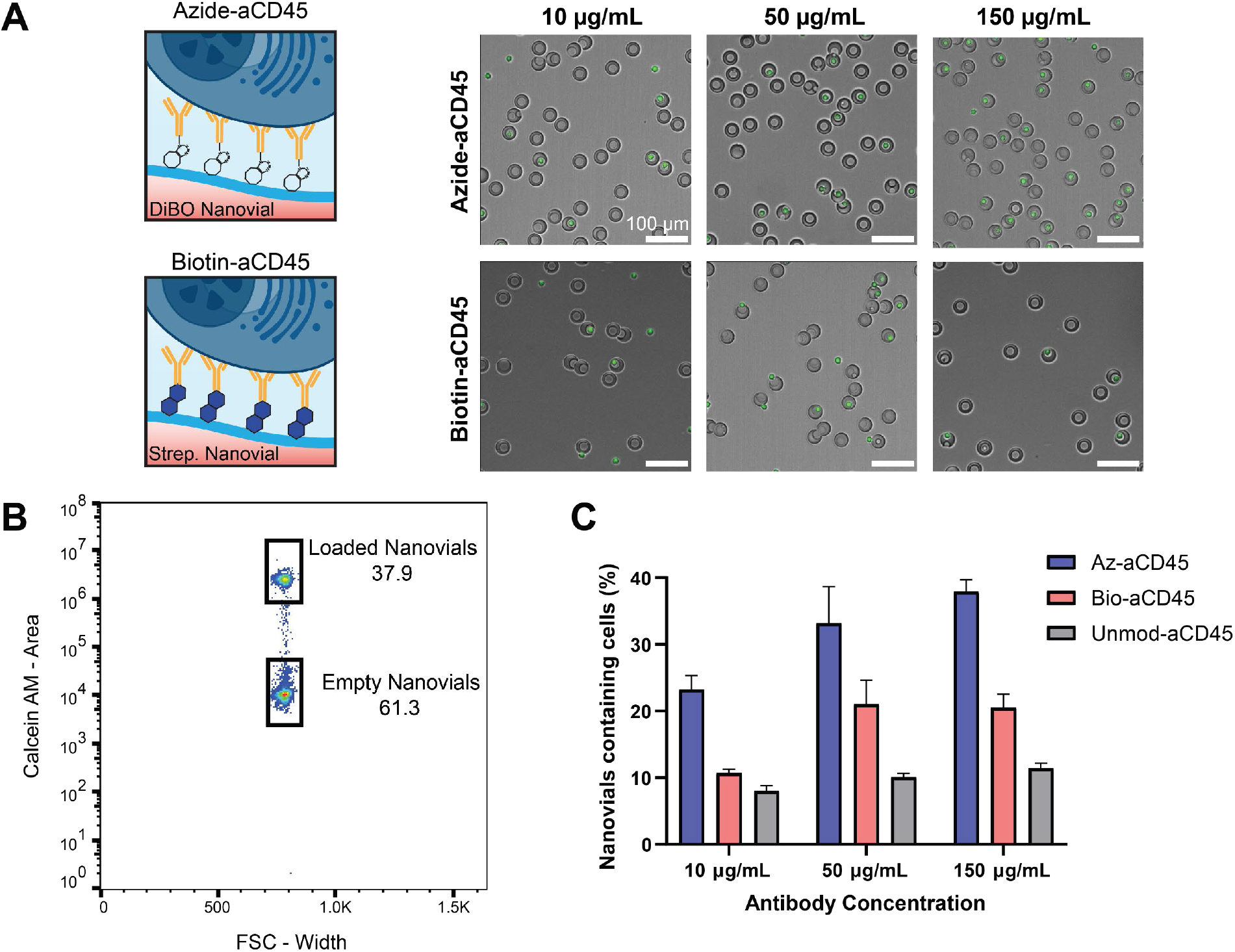
Azido-modified antibodies promote live hybridoma attachment to nanovials. A) Schematic of cell loading assay using DiBO-conjugated or biotin-conjugated nanovials and fluorescence microscopy of hybridoma cells (green) loading into nanovials with biotinylated aCD45 or azido-modified aCD45 capture antibodies. Scale bars are 100 microns. B) Flow cytometry gating results. High calcein AM area signal corresponds to nanovials loaded with a live cell, whereas low signal corresponds to empty nanovials. C) Cell loading efficiency is measured by counting the percentage of nanovials that contain a cell. Various concentrations of biotinylated, azido-modified, and unconjugated aCD45 antibodies are compared for relative loading rates.

### Orthogonal chemistries reduce crosstalk and improve hybridoma enrichment and purity

To test the hypothesis that adding capture antigens following cell loading would lead to crosstalk reduction, we performed a mock hybridoma campaign. We used a mixed population of hybridoma clones composed of 20% HyHEL5 (stained with calcein AM) and 80% 9E10 hybridomas (Fig 4A). The HyHEL5 antibody binds to the hen egg lysozyme (HEL) antigen, while the 9E10 antibody binds to the myc polypeptide tag. We prepared mixtures of the two hybridoma populations to directly compare two workflows. The first workflow used orthogonal chemistry-based screening, where the capture antigen was added to nanovials after cell loading. The second workflow employed biotin-only chemistry, where the capture antigen was added simultaneously with the cell capture antibody and was present during cell loading. In both assays we used biotinylated HEL as our secretion capture antigen. Only antibodies secreted from the HyHEL5 hybridomas will bind to the HEL antigen on the nanovial surface and any secretion capture on an unloaded nanovial or a nanovial loaded with a 9E10 cell will be the result of crosstalk from nearby HyHEL5 clones. We used azido-modified and biotinylated anti-mouse CD45 for cell capture for the orthogonal chemistry and biotin only workflows, respectively. Secreted antibodies were detected on the nanovials using DyLight 650 anti-mouse IgG antibodies. By plotting cell-loaded nanovials for DyLight 650 area (secretion) signal (Fig 4B), we sorted the top 2.5%, 5%, 10%, and 20% of DyLight 650 signal events and measured the sorted population purity (Fig 4C & 4D) by counting the number of calcein AM positive cells (HyHEL5 events) compared to the total number of sorted, cell-loaded nanovials (HyHEL5 + 9E10 events). We found that across every gating stringency the orthogonal chemistry workflow yielded better purity than the biotin only system. The top 2.5% of DyLight 650 signal events led to purities of 96.1% and 67.4% using orthogonal chemistries and biotin only, respectively (Fig 4C). Fluorescence microscopy of the nanovials prior to sorting shows appreciable background fluorescence on nanovials that are empty or contain unstained (9E10) cells for the biotin-only workflow, whereas fluorescence appears localized on nanovials containing calcein AM positive cells for the orthogonal chemistry workflow (top panel, Fig 4D). Post-sort imaging (bottom panel, Fig 4D) revealed a larger fraction nanovials containing calcein AM positive cells using the orthogonal chemistry workflow. This was driven by a rightward shift of the median anti-IgG detection antibody area signal (right panel, Fig. 4B) in the biotin only workflow. It can also be visualized by plotting the anti-IgG secretion detection height and area for empty nanovials and nanovials loaded with HyHEL5 cells (SI Fig 2A). The median secretion signal intensity is 41.8 times higher for HyHEL5-loaded nanovials than empty nanovials when using orthogonal chemistries, but only 3.23 times higher when using biotin only (SI Fig 2B). Reflecting the reduced crosstalk to empty nanovials and nanovials containing non-target cells, analysis of cell-loaded nanovials using orthogonal chemistries revealed a separate distinct population of high secretion signal events that represented approximately 10% of the total cell-loaded population (Fig 4B).

**Figure 4.**
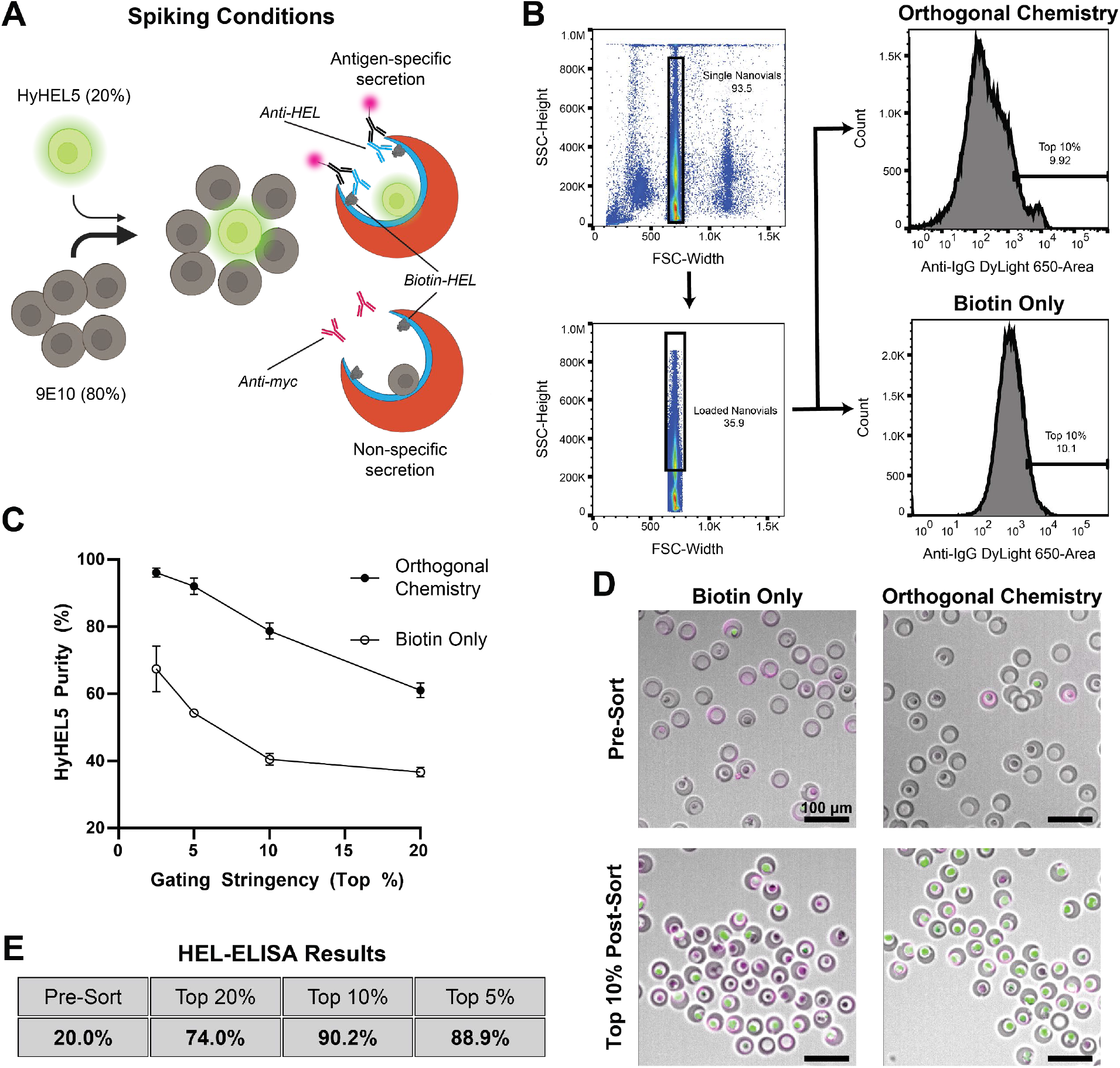
Orthogonal chemistry assay improves hybridoma screening performance. A) Overview of spiking conditions. Antigen-specific HyHEL5 hybridomas are spiked into non-antigen-specific 9E10 background population at a ratio of 20%/80%. B) Gating strategy for hybridoma enrichment. Single nanovial events are gated by plotting side scatter height against forward scatter width. Nanovials loaded with cells are gated using high side scatter height signal. This population is plotted for anti-IgG DyLight 650 area signal, which is used for sorting. C) Graph of HyHEL5 purity vs. gating stringency for orthogonal (filled circles) and biotin (open circles) chemistries. HyHEL5 purity is measured by counting the ratio of nanovials loaded with calcein-AM positive/negative cells after sorting exclusively for anti-IgG signal, using the top 2.5%, 5%, 10%, and 20% from the histogram in B. D) Fluorescence microscopy and overlaid brightfield images of orthogonal chemistry and biotin only nanovials loaded with the mixed hybridoma pools and stained for antigen-specific IgG secretion. Pre-Sort are samples prior to FACS sorting, while Top 10% Post-Sort reflects enriched samples following gating on highest 10% of anti-IgG signal. Scale bars are 100 microns. E) A table of the ELISA results for percent of supernatants that were reactive to HEL for initial pre-sort populations and following re-culture of sorted clones for increasing gating thresholds (Top 20% to Top 5%).

To demonstrate compatibility with typical hybridoma analysis and downstream processing workflows, we also performed single-cell sorts from the top 5%, 10%, and 20% of secretors for regrowth and mAb characterization. Single cells expanded from nanovials decorated with either biotinylated or azido-modified anti-CD45 antibodies (SI Fig 3A), doubling at expected rates. Culturing HyHEL5 cells in contact with either multi-reactive or standard nanovials does not affect cell viability (SI Fig 3B). Viability and regrowth are essential to establish new hybridoma colonies from selected pools. Finally, after allowing hybridomas to expand for two weeks after sorting, we performed an ELISA on the supernatants to characterize the purity and function of the clones selected using the orthogonal assay. We found that mAbs from the supernatants of selected clones bound to HEL antigen with the fraction of clones secreting HEL-specific mAbs in agreement with levels observed from measuring the fraction of calcein AM-positive clones (Fig 4E).

## Conclusions

Lab-on-a-particle technologies promise to increase access to cutting-edge molecular and cellular assays. Here, we demonstrated that by adding orthogonally-reactive groups to nanovials we could add functionality to a single-cell secretion assay by decoupling earlier cell loading and background cell removal steps in time from secretion capture steps, ultimately increasing purity of selected antigen-specific ASCs in an antibody discovery workflow. The increased purity was driven by a sharp reduction in crosstalk when adding secretion capture antibodies subsequent to all cell loading and filtration steps, enabled by using orthogonal chemistries on the nanovial surface. This methodology only requires an antibody functionalization (azide conjugation) step to be conducted before the assay and can be easily integrated with other types of nanovial-based screening assays, or other lab-on-a-particle formats for molecular and cellular assays.^24^ This approach is particularly beneficial to applications where cells are secreting at high levels or a large fraction of cells are actively secreting a target protein. For example, pre-enriched hybridoma pools, producer cells (such as CHO and HEK293 cell) screens, or analysis of IgG production rates by plasma cells, could all benefit from reduced crosstalk.

Covalent modification of nanovials may also provide unique advantages, such as cell loading efficiency, barcoding, and improved assay stability. Cell loading based on click chemistry-based conjugation was improved over biotin-streptavidin chemistry, and should be further investigated. The results highlight that cell loading rates can be improved substantially through changes in cell capture antibody presentation or density. Covalent and stable modification with capture antibodies could also reduce the number of steps to use nanovials, following manufacture and functionalization. Azido-modified oligonucleotide sequences can be used for barcoding particles for multiplexed assays, where covalent attachment provides improved stability over biotin-streptavidin linkages.^25^ In addition, molecular assays using samples rich in biotin or detection antibodies that have biotin/streptavidin conjugates would benefit from click chemistry-based linkages since the presence of free biotin would not obstruct any conjugation steps and downstream results. Similarly, assays where cells must be cultured in high concentrations of biotin, such as certain hybridoma and Chinese hamster ovary (CHO) strains, click chemistry could be beneficial.

More broadly, the use of multi-reactive nanovial surfaces enables new opportunities in nanovial assay development. It is possible to tune the relative amount of biotin and DiBO on the nanovial cavity, which could be used to conjugate precise ratios of different capture antibodies. This approach would be useful for simultaneously detecting high-secreted and low-secreted proteins. In addition, the temporal secretion profiles of molecules, such as two different cytokines, could be captured and integrated over different time periods by attaching orthogonally-functionalized capture antibodies at sequential starting points during a secretion assay. Such dynamic studies could help understand how cytokine secretion is sequenced in time by a cell to achieve a particular function. Another potential application is in two-cell interaction assays, where different surface chemistries could be used to load two different cell types, potentially improving co-loading efficiencies for both.^21^ Using separate chemistries, it could also be possible to separate T cell loading from T cell activation, adding peptide major histocompatibility complex monomers following loading with anti-CD45 capture antibodies, for example, which could help reduce crosstalk in T cell assays.^26^

An enhanced ability to rapidly identify viable, antigen-specific hybridoma clones from a mixed population can be broadly applied to many areas of research. Providing new tools, like multi-reactive nanovials, that can be easily integrated with standard laboratory processes and screened at a rate of hundreds of cells per second will enable researchers to conduct high-throughput hybridoma campaigns using standard FACS instruments. These workflows can also be translated to screen primary B cells and other antibody-secreting cell types, which will enable exploration of a much greater diversity of antibody clones and lead to improved therapeutics and diagnostic tools. Ultimately, continued innovation in lab-on-a-particle tools will yield more sophisticated functional understanding of diverse cellular populations and the enrichment of useful rare cells and cell lines.

## Materials and methods

### Culture of hybridoma cells

HyHEL5 hybridomas were provided by Richard Wilson from the University of Houston Department of Biology and Biochemistry. 9E10 hybridoma cells were purchased from ATCC. Both cell lines were cultured in IMDM cell culture media (Thermo Fisher 12440053) supplemented with 1% antibiotic-antimycotic (Thermo Fisher 15240062) and 10% fetal bovine serum (Thermo Fisher A5669701). Media was sterile filtered using 0.22 μm, 500 mL Stericups (Thermo Fisher S2GPU05RE). Cells were rapidly thawed from liquid nitrogen at 1 × 10^5^ cells per mL and passaged three times per week. Cells were maintained in a sterile incubator at 37°C and 5% CO_2_ and passaged up to P20 before replacement with a fresh vial.

### Nanovial particle fabrication

Nanovial fabrication has been described extensively in previous publications.^19,20^ Briefly, a polydimethylsiloxane (PDMS) flow focusing droplet generator is used to fabricate nanovials. A PEG solution composed of 27.5% w/v 4-arm, 5 kDa PEG-acrylate (Advanced BioChemicals, 4AP0902-1g) and 4% w/v lithium phenyl-2,4,5-trimethylbenzoylphosphinate (Sigma, 900889) in phosphate buffered saline (PBS) (Thermo Fisher, 14190250) was co-flowed with a gelatin solution composed of 20% w/v cold water fish skin gelatin (Sigma, G7041100G) in sterile-filtered DI water and an oil phase composed of 0.5% w/w Pico-Surf (Sphere Fluidics, C024), in sterile-filtered Novec^TM^ 7500 Engineered Fluid (3M, 7100134816). The PEG, gelatin, and oil solutions are mounted onto individual syringe pumps and perfused at 1, 1, and 15 μL per minute, respectively. Once stable droplets are formed and the gelatin phase separates from the PEG phase, the PEG phase is crosslinked using a UV lamp through a 10× microscope objective. Particles and excess oil are collected in a tube and a layer of PBS is added to the emulsion. Excess Pico-Surf is removed by washing the emulsion three times with sterile filtered Novec^TM^ 7500. Emulsions are broken using a 20% v/v solution of perfluoro-1-octanol (PFO) (Sigma, 370533) in Novec^TM^ 7500. Excess oil is aspirated and washed three times with hexane to remove any residual oil. Excess hexane is aspirated and the particles are washed three times with sterile-filtered 70% v/v ethanol to remove any excess hexane. Particles are filtered with a 70 μm filter (Stem Cell Technologies, 27216) to remove any aggregated nanovials. Nanovials are incubated overnight at 4°C in ethanol for sterilization. Finally, nanovials are washed three times in Pluronic wash buffer composed of 0.05% w/v Pluronic F-127, (Sigma, P2443), 1% v/v antibiotic-antimycotic, and 0.5% w/v bovine serum albumin (BSA) (Sigma, A7906) and stored in a 15 mL conical tube at 4°C for long-term storage.

### Nanovial surface functionalization

A 20 mM solution of dibenzocyclooctyne-PEG4-N-hydroxysuccinimidyl (DiBO-PEG-NHS) (Sigma, 764019) was prepared by dissolving 1 mg of DiBO-PEG-NHS paste in 77 μL of DMSO. A 20 mM solution of biotin-NHS (ApexBio, A8001) was made in DI H2O. For nanovials conjugated with both chemistries, a final solution of 1 mM DiBO-PEG-NHS and 10 mM biotin-NHS was made in DI water. A solution of 10 mM biotin-NHS in DI H2O was used for nanovials with a single chemistry. Each nanovial sample was incubated with 20 μL of each solution per 100,000 nanovials overnight with agitation at room temperature. The next day nanovials were washed three times with Pluronic washing buffer to remove excess DiBO-PEG-NHS and biotin-NHS. Nanovials were used for assays within one week of functionalization.

### Preparation of modified capture proteins

250 μg of anti-mouse CD45 antibody (Thermo Fisher, 50-115-49) was azido-modified using a SiteClick^TM^ Antibody Azido Modification Kit (Thermo Fisher, S20026) following the manufacturer’s protocol. Following conjugation, the antibody was purified on a column and concentration is measured using a spectrophotometer. Antibodies are stored at 4°C. Recombinant HEL (Aviva Systems Biology, OORA00201) was biotinylated using an EZ-Link^TM^ Sulfo-NHS-LC-Biotinylation Kit (Thermo Fisher, 21435) following the manufacturer’s protocol. Following purification, concentration is measured with a spectrophotometer and the protein is aliquoted and stored at -20°C. Azido-modified antibody conjugation was detected using an Alexa Fluor 647 anti-IgG antibody and image analysis was performed using ImageJ.

### Single-cell antibody capture assay

HyHEL5 and 9E10 cells were cultured and assessed for viability >95% before proceeding with any secretion experiments. 1 × 10^6^ HyHEL5 cells were stained with 1 μg per mL of calcein AM for 40 minutes at 37°C. HyHEL5 cells were washed three times with Pluronic washing buffer to remove excess dye. 9E10 and HyHEL5 cells were counted again and mixed into a ratio of 4:1 prior to both assay formats. For all assay steps microcentrifuge tubes, pipette tips, conical tubes, and other labware were pre-coated with Pluronic washing buffer to prevent nanovial accumulation.

### Biotin-only assay format

1.8 × 10^5^ 35 μm biotin-only nanovials were incubated with 75 μL of 300 μg/mL streptavidin (Thermo Fisher, 434302) in a 1.5 mL microcentrifuge tube for 30 minutes at room temperature, then washed three times with Pluronic washing buffer. Nanovials were incubated with a 50 μL mixture of 50 μg/mL biotinylated anti-mouse CD45 antibody (Thermo Fisher, 13045182) and 15 μg/mL biotinylated HEL antigen for 60 minutes at room temperature, then washed three times. 3.0 × 10^5^ cells from the mixed hybridoma pool were mixed with the functionalized nanovials in the well of a 12 well plate and placed on a rocker in a 37°C incubator for 60 minutes. Every 20 minutes the cells and nanovials were mixed by pipetting to promote cell loading. Following loading, unbound cells were removed using a 20 μm CellTrics cell strainer (Thermo Fisher, NC9699018) and nanovials with and without cells were reverse strained into a well of a 6 well plate with IMDM media. Cells were allowed to incubate without agitation at 37°C for 30 minutes to enable secretion accumulation. After washing two times with Pluronic washing buffer, nanovials were incubated with 50 μL of 25 μg/mL anti-mouse IgG Fc DyLight 650 antibody (Abcam, ab98715). After incubating for 30 minutes with rotation in the dark, nanovials were washed two more times and then transferred to FACS tubes for flow cytometry.

### Orthogonal chemistry assay format

1.8 × 10^5^ 35 μm biotin and DiBO-functionalized nanovials were incubated with a 50 μL mixture of 50 μg/mL azido-modified anti-mouse CD45 antibody for 60 minutes at room temperature, then washed three times. 3.0 × 10^5^ cells from the mixed hybridoma pool were loaded onto nanovials as described above and unbound cells were removed via reverse straining. Cell-loaded nanovials were mixed with 75 μL of 300 μg/mL streptavidin for 30 minutes, then washed three times, followed by 50 μL of 15 μg/mL biotinylated HEL for 60 minutes and three additional washes, all at room temperature. Cell-loaded nanovials were transferred to a well of a 6 well plate and allowed to incubate for 30 minutes at 37°C for secretion accumulation. Cells were similarly washed, stained with anti-mouse IgG Fc DyLight 650 antibody, and transferred to FACS tubes as above. Single-sorted cells were allowed to grow for 2 weeks before supernatant collection. ELISA plates were conjugated with 2 μg/mL of recombinant HEL antigen. After supernatant samples were added, ELISA plates were washed three times and conjugated with an HRP-conjugated, anti-mouse antibody (Abcam, ab205719). After addition of o-phenylenediamine, well absorbance levels were measured using a plate reader. Any samples greater than three standard deviations above filtered IMDM media were classified as HyHEL5 positive.

### Flow cytometry

Flow cytometric analysis and sorting was performed using a Sony SH800S Cell Sorter (Sony Biotechnology) for both assay formats. The DyLight 650 and Alexa 647 streptavidin fluorophores were excited using a 638 nm laser filtered through a 665/30 filter and calcein AM and FITC azide were excited using a 488 nm laser filtered through a 525/50 filter. For nanovial cell secretion assays, after gating for singlet nanovials, cell-loaded nanovials were gated by selecting nanovial events that had elevated side scatter height. For both assay formats, 100 nanovials were sorted into wells based on DyLight 650 area intensity with three replicates per gating stringency. Sorting was performed on single-cell sort mode with drop delay set to 14. Fluorescence microscopy was used to count the ratio of calcein AM positive cells in loaded nanovials.

## Acknowledgements

Flow cytometry was performed in the UCLA Jonsson Comprehensive Cancer Center. The work is partially supported by grants TLD1K132758: KUH-ART and CA256084 from the National Institutes of Health and grant 2023-332386 from the Chan Zuckerberg Initiative Donor Advised Fund (CZI DAF), an advised fund of the Silicon Valley Community Foundation. Y.N. is supported by postdoctoral fellowships from Japan Society for the Promotion of Science and Nakatani Foundation.

