## Supplemental Figures for "Multi-reactive hydrogel nanovials for temporal control of secretion capture from antibody-secreting cells"

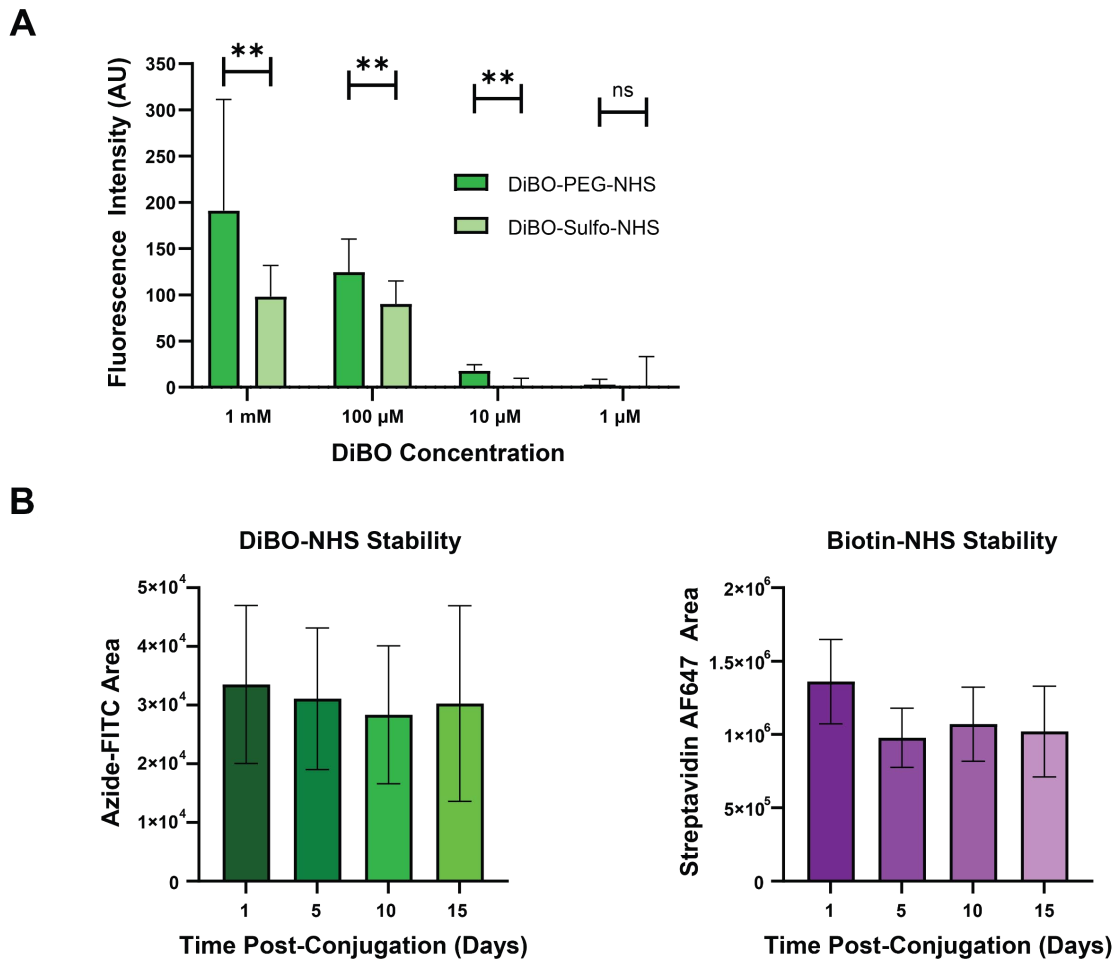


**SI Figure 1 DiBO surface modification and stability.** A) Image analysis of nanovials conjugated DiBO-PEG-NHS or DiBO-Sulfo-NHS and reacted with azide-FITC. ** Indicates p-value < 0.01 via two-way ANOVA t-test. B) Flow cytometry analysis of nanovials conjugated with 10 mM biotin-NHS and 1 mM DiBO-PEG-NHS and reacted with azide-FITC and streptavidin-Alexa Fluor 647. Biotin and DiBO conjugations were performed at different time points prior to fluorophore attachment and measurement.

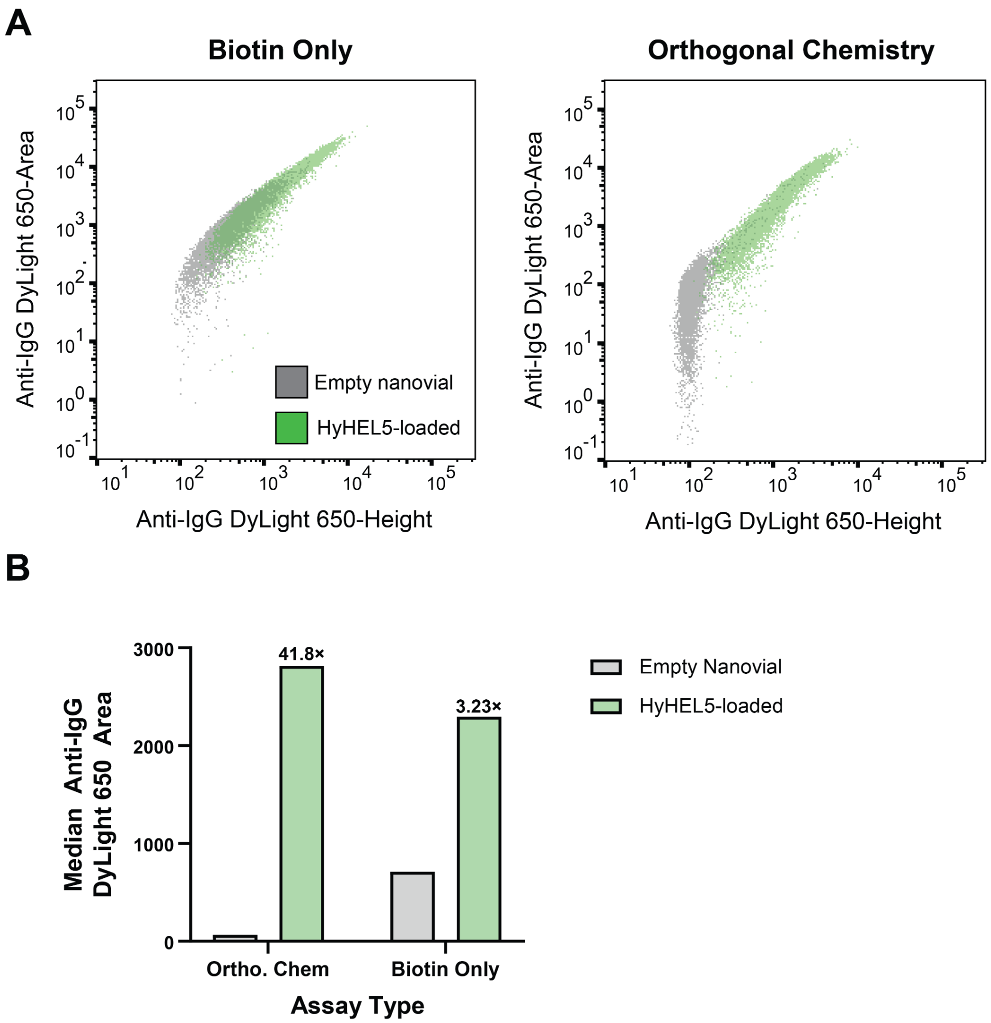


**SI Figure 2 Empty nanovial vs. HyHEL5-loaded secretion capture profiles.** A) Flow cytometry scatter plots of the anti-HEL specific mAb capture signal height and area for empty nanovials and nanovials loaded with HyHEL5 cells. Overlap between the populations is visible using the biotin only workflow, whereas there is much less using orthogonal chemistries. B) Median anti-IgG DyLight 650 secretion signal on HyHEL5-loaded and empty nanovials are plotted for the orthogonal (Ortho. Chem) and biotin only chemistries. The orthogonal chemistry workflow results in a nearly 42-fold difference in median secretion signal, compared to a 3-fold difference in the biotin only workflow.

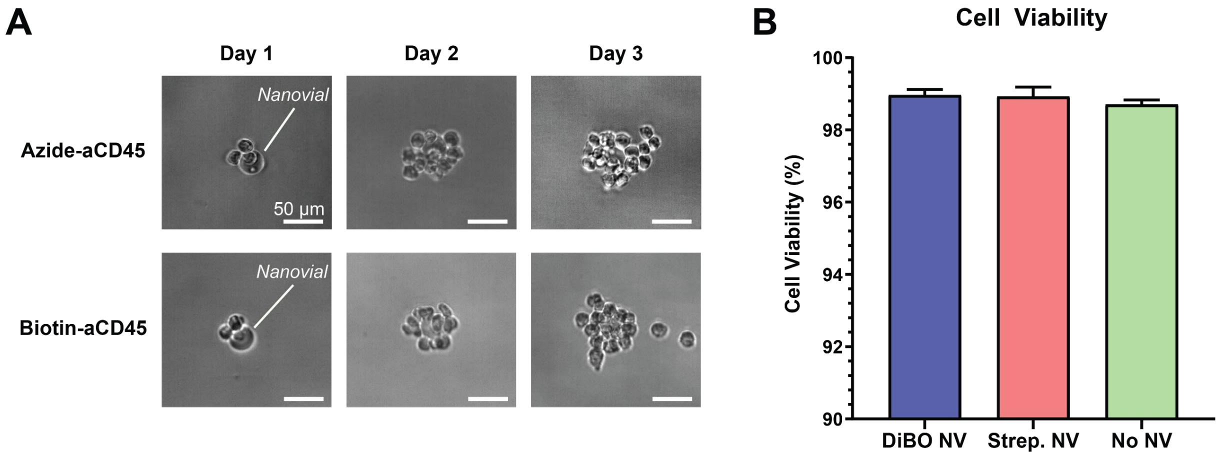


**SI Figure 3 Clonal expansion and cell viability unaffected by nanovial surface chemistry.**
A) Brightfield microscopy of HyHEL5-loaded nanovial sorts following multi-day incubations using azido-modified aCD45 or biotinylated aCD45 capture antibodies. Scale bars are 50 microns. B) Percentage of viable cells measured by calcein AM positive fluorescence following 24-hour incubation with nanovials conjugated with biotin and/or DiBO.
